# *Tamarixia citricola* Hansson and Guerrieri sp. nov. (Hymenoptera: Eulophidae): a new parasitoid of *Diaphorina citri* Kuwayamava (Hemiptera: Psyllidae) found during a classical biological control program in Cyprus

**DOI:** 10.1101/2025.06.16.660002

**Authors:** Alberto Urbaneja, Meritxell Perez-Hedo, Nicos Seraphides, Anthemis Melifronidou-Pantelidou, Stalo Giallouridou, Lia Markou, Menelaos C. Stavrinides, Chrysos Kaponas, Omar Ruiz-Rivero, David J.W. Morgan, Christer Hansson, Mark S. Hoddle, Alejandro Tena, Emilio Guerrieri

## Abstract

Asian citrus psyllid, *Diaphorina citri*, is a major global pest because it is the primary vector of *Candidatus* Liberibacter spp., the causal agents of huanglongbing (HLB), a lethal citrus disease. Following the detection of *D. citri* in Cyprus in 2023, the first record of this pest in the European Union, a classical biological control program targeting this pest was initiated in spring 2024 using the parasitoid *Tamarixia radiata* imported from California, USA. During field surveys in summer 2024, parasitized *D. citri* nymphs were found in orchards where no *T. radiata* releases had been made. These findings suggested the possible presence of native or unintentionally introduced parasitoids, or a rapid spread of *T. radiata* into new areas. To determine the identity of parasitoids associated with *D. citri* in Cyprus, an integrative approach was adopted combining field observations, molecular analyses of the COI gene, and morphological analyses. *Tamarixia radiata* recovered from Cyprus field sites matched reference sequences of parasitoids from California. However, other specimens were genetically and morphologically distinct and represented a new species. The new species is described here as *Tamarixia citricola* Hansson & Guerrieri sp. nov. Taxonomic diagnoses and characters for separating both *Tamarixia* species associated with *D. citri* are provided. Results presented here indicate the coexistence of both *T. radiata* (introduced) and *T. citricola* (likely autochthonous) in Cyprus citrus orchards. This finding has important implications for future biological control strategies and quarantine measures for *D. citri* in the Mediterranean basin.

## 1. Introduction

Huanglongbing (HLB), also known as citrus greening disease, is currently one of the most devastating threats to global citriculture. The disease is caused by phloem-restricted bacteria of the genus *Candidatus* Liberibacter, primarily *C.* L. asiaticus, *C.* L. africanus, and *C.* L. americanus, which are mainly transmitted by two psyllid vectors: Asian citrus psyllid (ACP), *Diaphorina citri* Kuwayama (Hemiptera: Psyllidae), and African citrus psyllid *Trioza erytreae* Del Guercio (Hemiptera: Triozidae) (Pérez-Hedo et al., 2025b, 2025a). Of these two psyllid species, *D. citri* is recognized as the most efficient and widely distributed vector, particularly of *C.* L. asiaticus (*C*Las), the most aggressive and globally prevalent bacteria causing HLB (Qureshi and Stansly, 2020). *C*Las spp. cause severe symptoms in citrus trees, including chlorosis, misshapen fruits, significant yield decline, and eventual tree death, leading to major economic losses and severe restrictions on international trade (Bové, 2014). In the last decades, *D. citri* has rapidly spread, invading citrus-growing regions across the Americas, Asia, and Africa, and more recently, the eastern Mediterranean basin (Pérez-Hedo et al., 2025a). In August 2023, *D. citri* was officially detected for the first time within the European Union (EPPO Global Database, 2023), in a citrus orchard located in the Phassouri area of Limassol District, Cyprus (Melifronidou-Pantelidou et al., 2025). This discovery in Cyprus followed a previous report of *D. citri* in Israel in 2021 (EPPO Global Database, 2022), demonstrating the progressive spread of the pest across eastern Mediterranean areas.

The detection of *D. citri* in Cyprus triggered an immediate and coordinated phytosanitary response led by the National Plant Protection Organization (NPPO) of Cyprus. This response included comprehensive island-wide surveillance, the establishment of demarcated *D. citri*-infested and *D. citri*-free buffer zones, enforced restrictions on the movement of citrus, and insecticidal treatments (Melifronidou-Pantelidou et al., 2025). However, due to the widespread distribution of citrus plants in commercial orchards, private gardens, recreational areas, and commercial spaces, chemical control measures alone were deemed insufficient to achieve a significant reduction of *D. citri* densities and rates of spread. Consequently, a classical biological control program was initiated by the Cyprus NPPO in collaboration with the Agricultural Research Institute of Cyprus, the Valencian Institute of Agricultural Research (IVIA, Spain), the University of California, Riverside (UCR), and the California Department of Food and Agriculture (CDFA). This program involved the importation, quarantine and mass rearing of the parasitoid *Tamarixia radiata* (Waterston) (Hymenoptera: Eulophidae) from California (USA) for targeted releases against *D. citri* populations. The California population of *T. radiata* was originally sourced from Punjab Pakistan and used in a classical biological control program against *D. citri* in California (Hoddle and Pandey, 2014).

*Tamarixia radiata* is a solitary ectoparasitoid, native to the Indian subcontinent that specifically parasitizes fourth and fifth-instar nymphs of *D. citri* (Hoddle and Pandey 2014; Hoddle et al., 2022). This parasitoid has been widely recognized for its potential in classical biological control programs due to its high host specificity, short generation time, high rates of parasitism, and host feeding behaviour. As a result, *T. radiata* has been intentionally introduced or reintroduced into multiple regions invaded by *D. citri*, including Florida, Brazil, and Mexico (Qureshi and Stansly, 2020).

In California, *T. radiata* was introduced in 2011 as part of a coordinated classical biological control initiative, following exploratory surveys and collections conducted in the Punjab region of Pakistan, an area sharing a high climatic similarity to California’s citrus production zones (Hoddle and Hoddle, 2013; Hoddle and Pandey 2014). The parasitoid quickly established in urban citrus landscapes and demonstrated high efficacy, with field studies reporting population reductions of *D. citri* up to 75% following releases (Kistner et al., 2016a; Milosavljević et al. 2021). Parasitism levels exceeded 60% at times, and the biological control effort was associated with a notable slowing of *C*Las spread in urban environments because of significant reductions of vector densities (Milosavljević et al. 2021; Hoddle et al., 2022).

Given its successful establishment and performance in California and the proven effectiveness of *T. radiata* in this Mediterranean-like climate, Cypriot authorities selected, imported, and mass-reared under controlled conditions the same Californian strain as the cornerstone of the Cypriot biological control campaign targeting *D. citri*. Releases of *T. radiata* in Cyprus began in April 2024 at four citrus-growing locations (Melifronidou-Pantelidou et al., 2025).

Here, results are reported of the first monitoring surveys conducted at the four *T. radiata* release sites and at 12 control sites where no parasitoid releases occurred. This experimental design was used to evaluate the rates of parasitoid dispersal into non-release areas and to assess the impacts of autochthonous natural enemies, especially naturally-occurring parasitoid species, on *D. citri* populations. Parasitoids recovered during field surveys were identified using an integrative taxonomic approach that combined COI-based molecular analyses, alfa morphology and biological data. This approach revealed the existence of two distinct parasitoid species of *Tamarixia*, with one being new to science.

## 2. Materials and methods

### 2.1. Mass rearing and field releases of *Tamarixia radiata* in Cyprus

In March 2024, the Agricultural Research Institute (ARI) of Cyprus established a *T. radiata* colony from 250 individuals provided by the California Department of Food and Agriculture (CDFA). These parasitoids originated from a colony maintained at the CDFA biological production facility in Riverside, California, which was originally founded from individuals collected in Pakistan by the University of California, Riverside (Hoddle and Hoddle, 2013; Hoddle and Pandey, 2014). To support the rearing of *D. citri* for parasitoid propagation, seeds of curry leaf (*Murraya koenigii* L., Rutaceae), a preferred host plant for *D. citri* that does not harbour *C.* Liberibacter spp., were provided by the CDFA in December 2023 and grown at ARI facilities. The use of plants grown from seed further ensured the absence of *C*Las infection.

The *T. radiata* colony was maintained on *D. citri* nymphs reared on *M. koenigii* plants under controlled environmental conditions (25 ± 1°C, 60–70% relative humidity, 14L:10D photoperiod). These rearing conditions enabled the continuous propagation of parasitoids which were used to maintain colony production and for field releases. Exemplar specimens from the established ARI colony were preserved in 96% ethanol and used as reference material for both molecular and morphological analyses in the present study. The first releases of *T. radiata* in Cyprus were made in April 2024 in the Phassouri area (Limassol district, 60 parasitoids), followed by additional releases in the districts of Nicosia (Potamia, April 2024, 60 parasitoids), and Ammochostos (Frenaros, April and June 2024, 40 and 60 parasitoids, respectively, and Avgorou, June 2024, 40 parasitoids) (Figure 1). All *T. radiata* releases were made in citrus orchards exhibiting active *D. citri* infestations at the time of parasitoid release. This biological control program in Cyprus is ongoing, with monitoring efforts in place to assess establishment, field dispersal, and parasitism levels of *T. radiata*.

**Figure 1.**
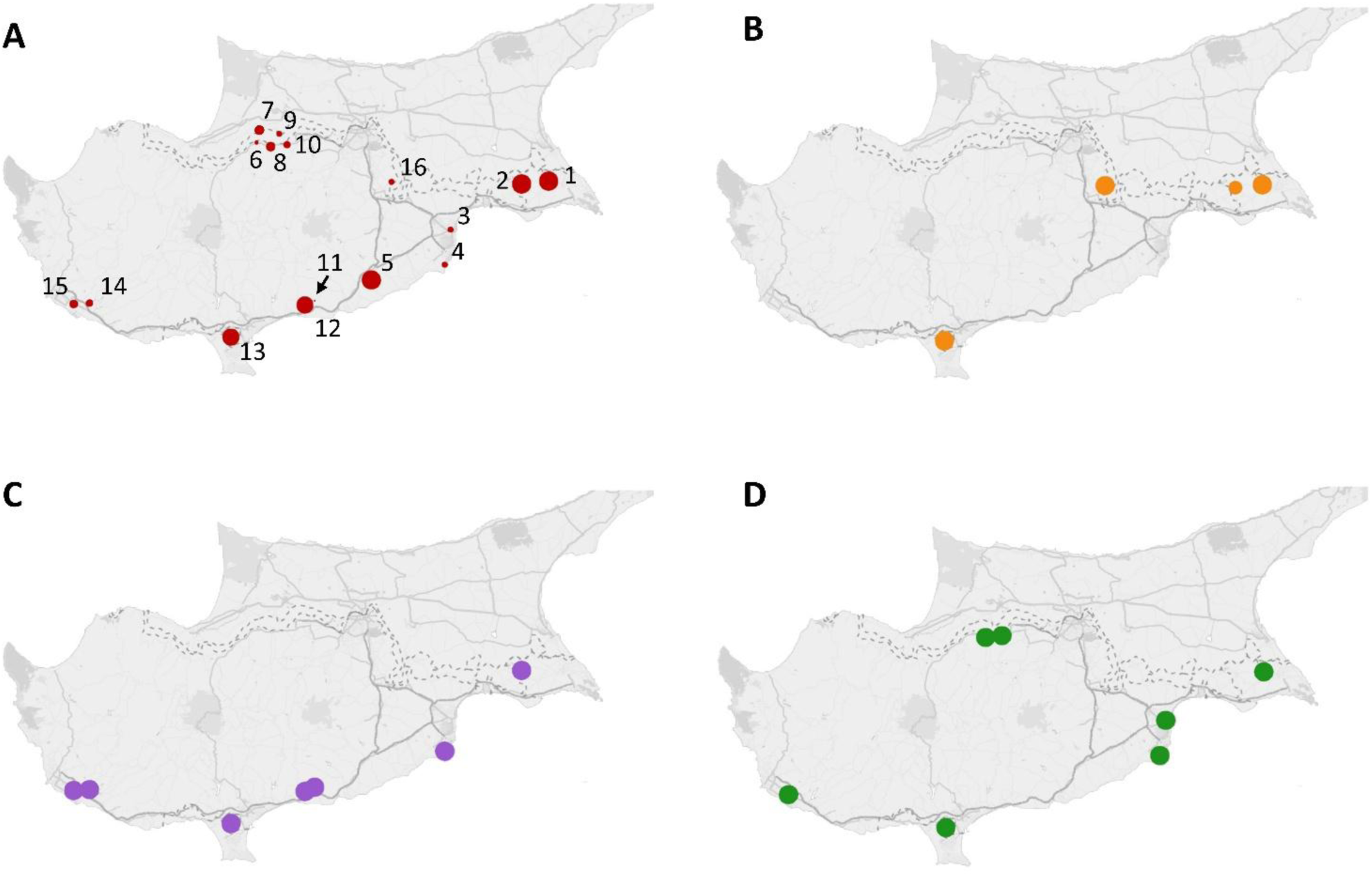
*Diaphorina citri* infestation and *Tamarixia radiata* presence across Cyprus in 2024. A) Density of *D. citri* on summer flush of citrus in July 2024, expressed as the percentage of infested shoots per site. The size of the red circles reflects infestation intensity, ranging from 1% to 100%, and can be compared with the exact values provided in Table 1. The sites are numbered as follows: (1) Frenaos, (2) Avgorou, (3) Meneou, (4) Meneou (2), (5) Agios Theodoros, (6) Peristerona, (7) Green Zone, (8) Peristerona 2, (9) Avlona, (10) Avlona (2), (11) Pyrgos, (12) Monagroulli, (13) Phassouri, (14) Anarita, (15) Acheleia, (16) Potamia. B) Locations where *T. radiata* was released in spring 2024. C) Recovery of *Tamarixia* spp. from spring flush samples. D) Recovery of *Tamarixia* spp. from summer flush samples.

**Table 1.**
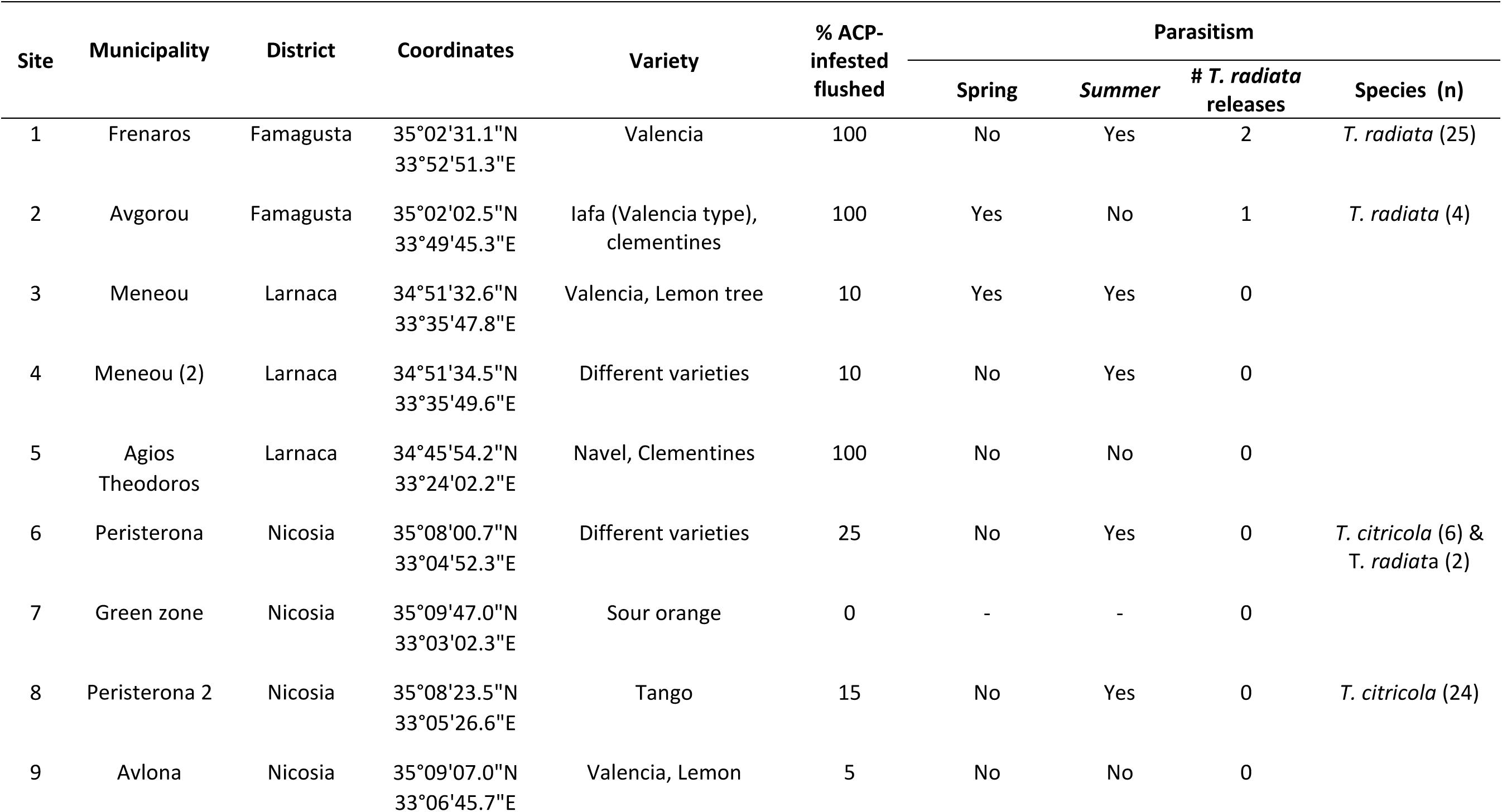
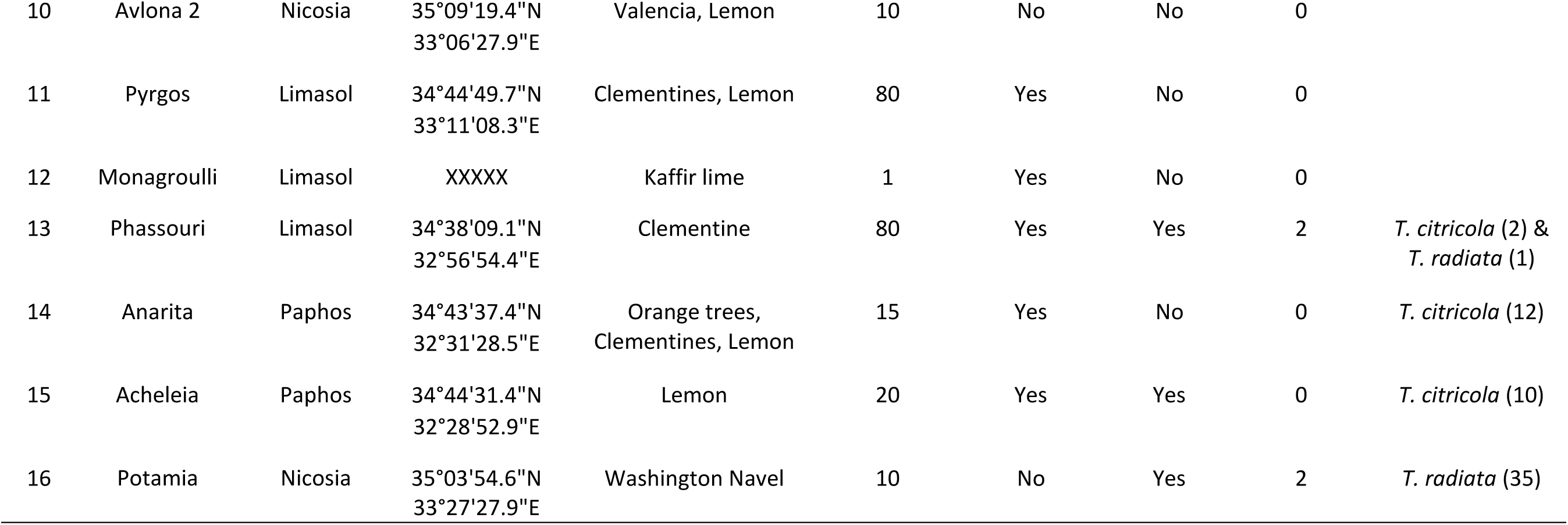
Sites surveyed for *Diaphorina citri* and parasitoid presence in Cyprus. The table describes the location (municipality, district, and GPS coordinates), citrus varieties present, percentage of flushing shoots infested by *D. citri* (ACP-infested flushed), seasonal presence of parasitoids in spring and/or summer, the number of releases of *Tamarixia radiata* conducted at each site, and the parasitoid species recovered, with the number of individuals collected in parentheses.

| Site | Municipality | District | Coordinates | Variety | % ACP-infested flushed | Parasitism |  |  |  |
| --- | --- | --- | --- | --- | --- | --- | --- | --- | --- |
|  |  |  |  |  |  | Spring | Summer | # *T. radiata* releases | Species (n) |
| 1 | Frenaros | Famagusta | 35°02'31.1"N<br>33°52'51.3"E | Valencia | 100 | No | Yes | 2 | *T. radiata* (25) |
| 2 | Avgorou | Famagusta | 35°02'02.5"N<br>33°49'45.3"E | Iafa (Valencia type),<br>clementines | 100 | Yes | No | 1 | *T. radiata* (4) |
| 3 | Meneou | Larnaca | 34°51'32.6"N<br>33°35'47.8"E | Valencia, Lemon tree | 10 | Yes | Yes | 0 |  |
| 4 | Meneou (2) | Larnaca | 34°51'34.5"N<br>33°35'49.6"E | Different varieties | 10 | No | Yes | 0 |  |
| 5 | Agios Theodoros | Larnaca | 34°45'54.2"N<br>33°24'02.2"E | Navel, Clementines | 100 | No | No | 0 |  |
| 6 | Peristerona | Nicosia | 35°08'00.7"N<br>33°04'52.3"E | Different varieties | 25 | No | Yes | 0 | *T. citricola* (6) &<br>*T. radiata* (2) |
| 7 | Green zone | Nicosia | 35°09'47.0"N<br>33°03'02.3"E | Sour orange | 0 | - | - | 0 |  |
| 8 | Peristerona 2 | Nicosia | 35°08'23.5"N<br>33°05'26.6"E | Tango | 15 | No | Yes | 0 | *T. citricola* (24) |
| 9 | Avlona | Nicosia | 35°09'07.0"N<br>33°06'45.7"E | Valencia, Lemon | 5 | No | No | 0 |  |
| 10 | Avlona 2 | Nicosia | 35°09'19.4"N<br>33°06'27.9"E | Valencia, Lemon | 10 | No | No | 0 |  |
| 11 | Pyrgos | Limasol | 34°44'49.7"N<br>33°11'08.3"E | Clementines, Lemon | 80 | Yes | No | 0 |  |
| 12 | Monagroulli | Limasol | XXXXX | Kaffir lime | 1 | Yes | No | 0 |  |
| 13 | Phassouri | Limasol | 34°38'09.1"N<br>32°56'54.4"E | Clementine | 80 | Yes | Yes | 2 | *T. citricola* (2) &<br>*T. radiata* (1) |
| 14 | Anarita | Paphos | 34°43'37.4"N<br>32°31'28.5"E | Orange trees,<br>Clementines, Lemon | 15 | Yes | No | 0 | *T. citricola* (12) |
| 15 | Acheleia | Paphos | 34°44'31.4"N<br>32°28'52.9"E | Lemon | 20 | Yes | Yes | 0 | *T. citricola* (10) |
| 16 | Potamia | Nicosia | 35°03'54.6"N<br>33°27'27.9"E | Washington Navel | 10 | No | Yes | 2 | *T. radiata* (35) |

### 2.2. Field surveys and sample collection

To assess the establishment and spread of *T. radiata* across Cyprus, field surveys were conducted between July 2 and 5, 2024. A total of 16 citrus orchards (Table 1), distributed across the main citrus-producing districts of Cyprus, were selected for evaluation. In four orchards *T. radiata* was released during spring 2024, while the remaining 12 served as non-release control sites.

In each orchard, the following parameters were recorded: citrus variety, presence of spring and summer flush (i.e., immature leaf and twig growth used by *D. citri* for oviposition and feeding by nymphs), and percentage of shoots infested by *D. citri*. The percentage of shoot infestation was determined in each orchard by the presence or absence of *D. citri* nymphs or adults on 20–30 shoots randomly selected on 5-8 trees that were randomly selected for sampling. *Diaphorina citri* nymphs on infested stems were examined for evidence of *T. radiata* parasitism. Parasitism was determined visually by detecting either parasitoid eggs, nymphs, or pupae under the ventral surfaces of fourth and fifth instar *D. citri* nymphs or circular exit holes in the anterior part of mummified nymphs (Kistner et al., 2016b, 2016a; Milosavljević et al., 2021).

Where possible, in orchards where active parasitism was observed, parasitized nymphs were collected for laboratory-based confirmation of parasitoid species identity. *Diaphorina citri* nymphs exhibiting signs of parasitism were sampled from eight study orchards. Individual field-collected parasitoids were preserved in 96% ethanol for molecular analyses.

### 2.3. Molecular analyses

Molecular analyses were conducted to enable unambiguous determination of the presence *T. radiata* in orchards and the geographic origin of parasitoid strain (i.e., from California or some other location). The correct identification of collected parasitoids provided baseline information on dispersal from release sites. To achieve this, a molecular barcoding approach using the mitochondrial cytochrome oxidase subunit I gene (COI) - formerly *COX1* gene - was conducted. Parasitoids were obtained from multiple locations across Cyprus and included both adult and pupal stages of eulophid parasitoids morphologically identified as *Tamarixia* sp.

Specifically, COI fragments 1A-1, 2A-1, and 11A-1 (were amplified from adult parasitoid specimens, while COI fragments 1B-1, 2B-1, and 5B-1 were obtained from *Tamarixia* sp. pupae collected from parasitized *D. citri* nymphs. In addition, a consensus COI fragment of *T. radiata* was generated from adults recovered from both field-collected specimens and from the laboratory colony established in Cyprus using parasitoids sourced from the California Department of Food and Agriculture (samples C1, C2, and C3). For comparative purposes, COI sequences of closely related species within the subfamily Tetrastichinae (Hymenoptera: Eulophidae), retrieved from GenBank, were incorporated into alignments.

Total mitogenomic DNA (mitgDNA) was extracted from each specimen using a modified salting-out protocol optimized for small arthropods (Monzó et al. 2011). A fragment of approximately 825 base pairs from the 3′ region of the COI gene was amplified from the specimens of *Tamarixia* sp. collected in the field using the primers C1-J-2183 (5’-CAACATTTATTTTGATTTTTTGG-3’) and TL2-N-3014 (5’-TTGCACTTTTCTGCCATTTTA-3’) (Simon et al., 1994). Given that most *T. radiata* COI gene sequences available in GenBank correspond to the 5’-end of its coding region (positions +106 to +676 relative to the translation start codon), the oligonucleotides LCO2 (5’-GGGCAACAAATCATAAAGATATTGG-3’) and HCOoutout (5’-GTAAATATATGRTGDGCTC-3’) (Folmer et al., 1994; Prendini, 2005, respectively), were also used as primers to amplify a COI fragment of 1487 bp from *T. radiata* specimens derived from the laboratory colony established in Cyprus. Polymerase chain reactions (PCR) were performed in a final volume of 20 μL using Platinum II Hot-Start Taq DNA Polymerase (ThermoFisher), 1× Platinum II buffer (containing 1.5 mM MgCl₂), 0.2 mM of each dNTP, and 0.2 μM of each primer, with 2 μL of mtDNA as template. The thermal cycling protocol consisted of an initial denaturation at 94°C for 2 minutes, followed by 10 cycles of 94°C for 15 seconds, 45°C for 15 seconds, and 68°C for 15 seconds, and then 40 additional cycles using the same parameters. PCR products were visualized on 1.5% agarose gels and purified using the High Pure PCR Cleanup Kit (Roche, Germany).

Purified amplicons were sequenced in both directions using the same primers. Forward and reverse reads were assembled into consensus sequences using Sequencher software (Gene Codes Corporation) and trimmed to remove primer sequences. The final high-quality consensus sequences (825 bp) were used as queries in BLASTn searches against the GenBank nucleotide database to assess species identity and similarity to known *Tamarixia* sp. species sequences. Alignments were performed using DNASTAR software, and the alignment of COI sequences used for phylogenetic analysis.

Phylogenetic reconstruction was performed using the Maximum Likelihood (ML) method implemented in MEGA12. The Tamura and Nei (1993) model of nucleotide substitutions was selected as the best-fit model. The initial tree for the heuristic search was chosen based on the best log-likelihood value obtained by comparing a Neighbor-Joining (NJ) tree, generated from a pairwise distance matrix using the same model, and a Maximum Parsimony (MP) tree. The MP analysis included 10 independent tree searches from randomly generated starting trees, and the shortest resulting tree was used for comparison. The tree with the highest log likelihood (−2,539.32) was selected as the final topology.

The robustness of the inferred phylogeny was assessed by bootstrap analysis with 1,000 replicates. The final dataset included 12 coding COI sequences from both adult-derived fragments (1A-1, 2A-1, 11A-1) and pupal-derived fragments (1B-1, 2B-1, 5B-1) of *Tamarixia* specimens collected from different locations in Cyprus. COI fragments from individuals of the *T. radiata* colony (samples C1, C2, C3) and field-collected adults were included as references. Additionally, COI sequences from related species within the subfamily Tetrastichinae (Hymenoptera: Eulophidae), retrieved from GenBank (Q310511, Q059861, NC_079567, MN123622, NC060368), were incorporated into the phylogenetic analysis. Codon positions included 1st, 2nd, 3rd, and non-coding sites, and all positions containing gaps or missing data were excluded, resulting in 825 aligned positions in the final dataset. All evolutionary analyses were conducted in MEGA12 using up to four parallel computing threads.

### 2.4. Morphological (alpha) characterization

*Tamarixia* specimens were prepared and mounted on card points and slide mounted. For card mounting, specimens freshly killed in 70% ethanol were transferred to 96% ethanol, then to absolute ethanol, and finally to hexamethyldisilazane (HMDS), where they were left until the HMDS completely evaporated. Once dried, specimens were mounted on cards using a water-soluble glue. For slide mounting, the protocol described by Noyes (1982) was followed. In brief, after detaching fore and hind wings, specimens were macerated in a 10% KOH solution on a hot plate for 3 minutes, immersed in acetic acid for 5 minutes, then dehydrated by placing in a series of increasing concentrations of ethanol from 70 %to 100%. Upon completion of this dehydration sequence, a drop of clove oil was added to the absolute ethanol and specimens were left until the ethanol evaporated. Specimens were removed from clove oil and mounted on glass slides in Canada balsam with the aid of a stereomicroscope (20x magnification). The morphological characterization of card and slide mounted *Tamarixia* specimens was initially performed using the key prepared by Graham(1991). Species identities were confirmed by comparing mounted specimens with type specimens and authoritatively identified material hosted at the Biological Museum in Lund and at the Natural History Museum in London.

Abbreviations used in the description of the new species:

- C1–C3: clavomeres 1–3 (in antennae),
- F1–F4; flagellomeres 1–4 (in antennae),
- Gt7: gastral tergite no. 7
- OOL: shortest distance between one posterior ocellus and eye
- POL: shortest distance between posterior ocelli
- T1–T4: tarsomeres 1–4 (of tarsi)

All measurements in the description of the new species are from card mounted specimens. These measurements do not always match measurements from slide mounted body parts because these might have been distorted when compressed between the slide glass and the cover slip.

## 3. Results

### 3.1. Field detection of parasitism by *Tamarixia* species

A total of 16 citrus orchards across five districts of Cyprus were surveyed for the presence of *D. citri* (Table 1). *Diaphorina citri* nymphs were detected in 15 out of 16 sites (93.7%), with infestation levels ranging from 1% to 100% of sampled flush (Table 1) (Figure 1A). Only one site, located in the Green Zone (site 7), showed no evidence of *D. citri*. Evidence of parasitism of *D. citri* nymphs was observed at 12 sites, based on abovementioned signs of parasitism. Among these, four sites (sites 1, 2, 13, and 16) had received intentional *T. radiata* releases (Figure 1B), while the remaining 8 sites (sites 3, 4, 6, 8, 11, 12, 14, and 15) had not received *T. radiata* (Figures 1C and 1D).

Adult *Tamarixia* spp. emerged successfully from field collected nymphs. A total of five sites yielded *Tamarixia* spp. adults, including *T. radiata* and an unknown *Tamarixia* species. Notably, adult *Tamarixia* emerged from samples collected in sites without any known history of parasitoid releases, confirming parasitism activity in those areas. Importantly, evidence of parasitism was detected during summer in several orchards located at more than 40 km from known release sites, including Acheleia, and Anarita (Table 1). This indicated early-season parasitism activity on *D. citri* in regions with no history of *T. radiata* introductions.

### 3.2. Molecular identification: COI analyses

A COI fragment of approximately 825 base pairs (bp) was successfully amplified and sequenced from all analyzed specimens, including individuals from the *T. radiata* laboratory colony and field-collected *Tamarixia* specimens recovered from parasitized *D. citri* nymphs collected across multiple field sites in Cyprus. The consensus sequences obtained from the laboratory colony (samples C1–C3) and field-collected *T. radiata* adults were identical and showed 99–100% similarity to *T. radiata* sequences deposited in GenBank (Supplementary Figure 1), including accessions from the USA (FJ152417 and FJ152420) and China (MZ558501), and 91% similarity to the *T. radiata* mitochondrial genome (MN123622). Importantly, molecular analyses confirmed recovery of *T. radiata* haplotypes introduced from California indicating that parasitoids deliberately introduced as classical biological control agents had established in Cyprus.

In contrast, COI sequences obtained from six field-collected *Tamarixia* specimens from Cyprus, three derived from adults 1A-1, 2A-1, 11A-1 (PV779178, PV779176, PV779180), and three from pupal stages 1B-1, 2B-1, 5B-1(PV779179, PV779177, PV779181), differed markedly from *T. radiata*, showing only 91–92% similarity to introduced *T. radiata* haplotypes and *T. triozae* (Q310511). These sequences were highly homogeneous with minimal intraspecific variation, and were consistently distinct from all known *Tamarixia* sequences available in public databases suggesting detection of a previously unknown species.

Alignment of the full 825 bp COI fragment (Figure 2) highlighted numerous nucleotide differences between the field-collected Cyprus specimens and *T. radiata*, as well as with other members of the subfamily Tetrastichinae (Eulophidae). Phylogenetic reconstruction based on this alignment, using the Maximum Likelihood method and the Tamura–Nei model, revealed the existence of two well-supported and genetically distinct clades (Figure 3). The first clade included all *T. radiata* samples from the Cyprus laboratory colony and field collected material, and reference sequences from GenBank. (MN132622). The second clade consisted exclusively of field-collected Cyprus specimens (both adult- and pupa-derived), which formed a monophyletic group strongly supported by bootstrap values ranging from 78 to 100.

**Figure 2.**
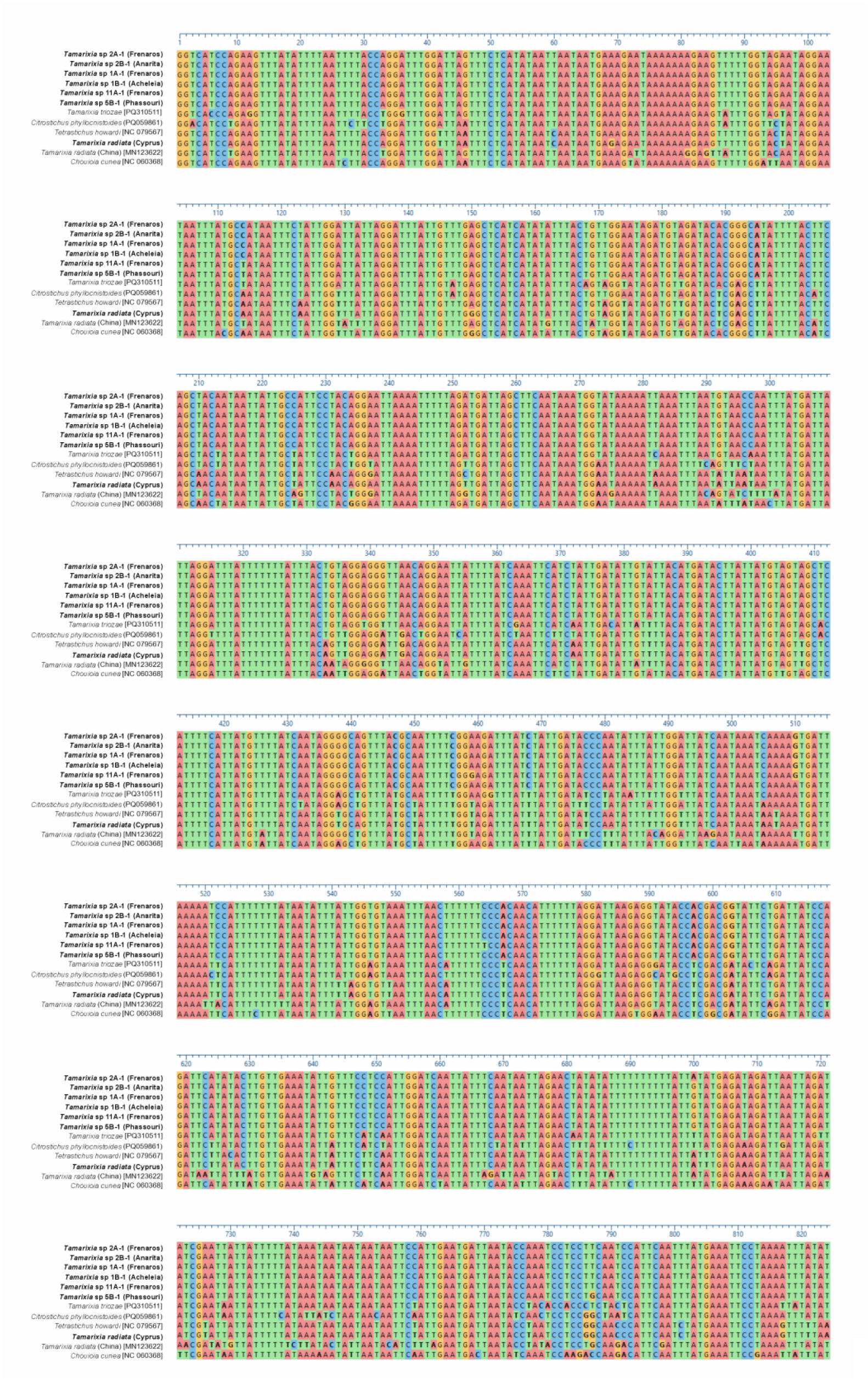
Multiple sequence alignment of mitochondrial COI gene fragments from *Tamarixia* spp. collected at different developmental stages (adults and pupae) and locations in Cyprus. A consensus sequence of *T. radiata* and COI sequences from related tetrastichine species obtained from GenBank were included for comparison. Variable sites across the 825 bp alignment are highlighted.

**Figure 3.**
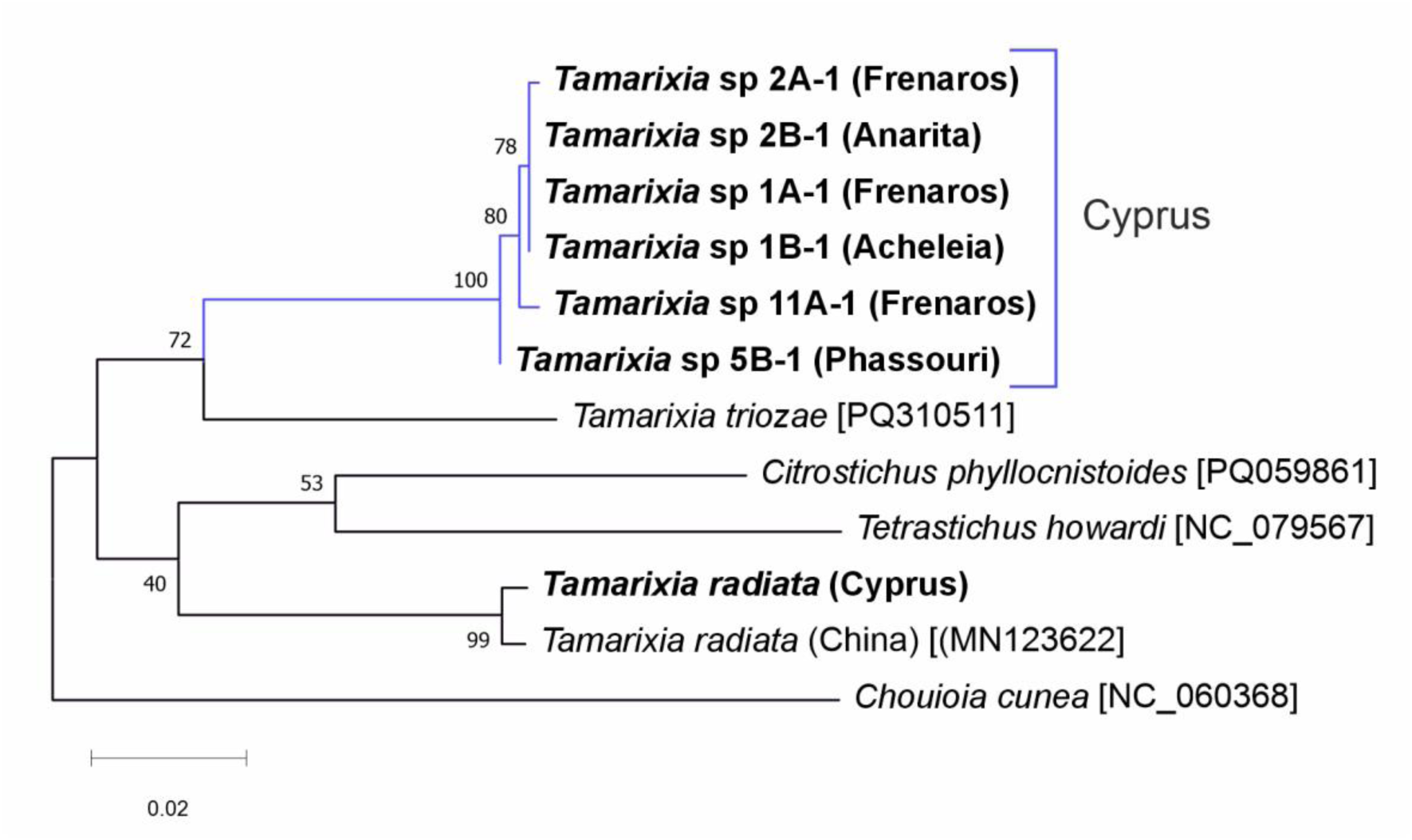
Maximum Likelihood phylogenetic tree based on COI gene sequences (825 bp) from *Tamarixia* spp. in Cyprus, including reference *T. radiata* sequences and closely related taxa from GenBank. Bootstrap values (1,000 replicates) are shown next to the branches. The evolutionary history was inferred using the Tamura–Nei model in MEGA12.

These findings provide robust molecular evidence for the presence of a genetically distinct lineage within the genus *Tamarixia* in Cyprus, which is clearly separated from *T. radiata* and other closely related species. This finding was interpreted as representing a new unidentified species, here named *Tamarixia citricola* sp. Nov. The corresponding COI sequences have been submitted to GenBank, with the following accession numbers: PV779178, PV779176, PV779180, PV779179, PV779177, PV779181.

### 3.3. Morphological identification and species description of *Tamarixia citricola* sp.nov. Hansson and Guerrieri

*Tamarixia citricola* sp.nov. Hansson and Guerrieri (Figure 4)

**Figure 4.**
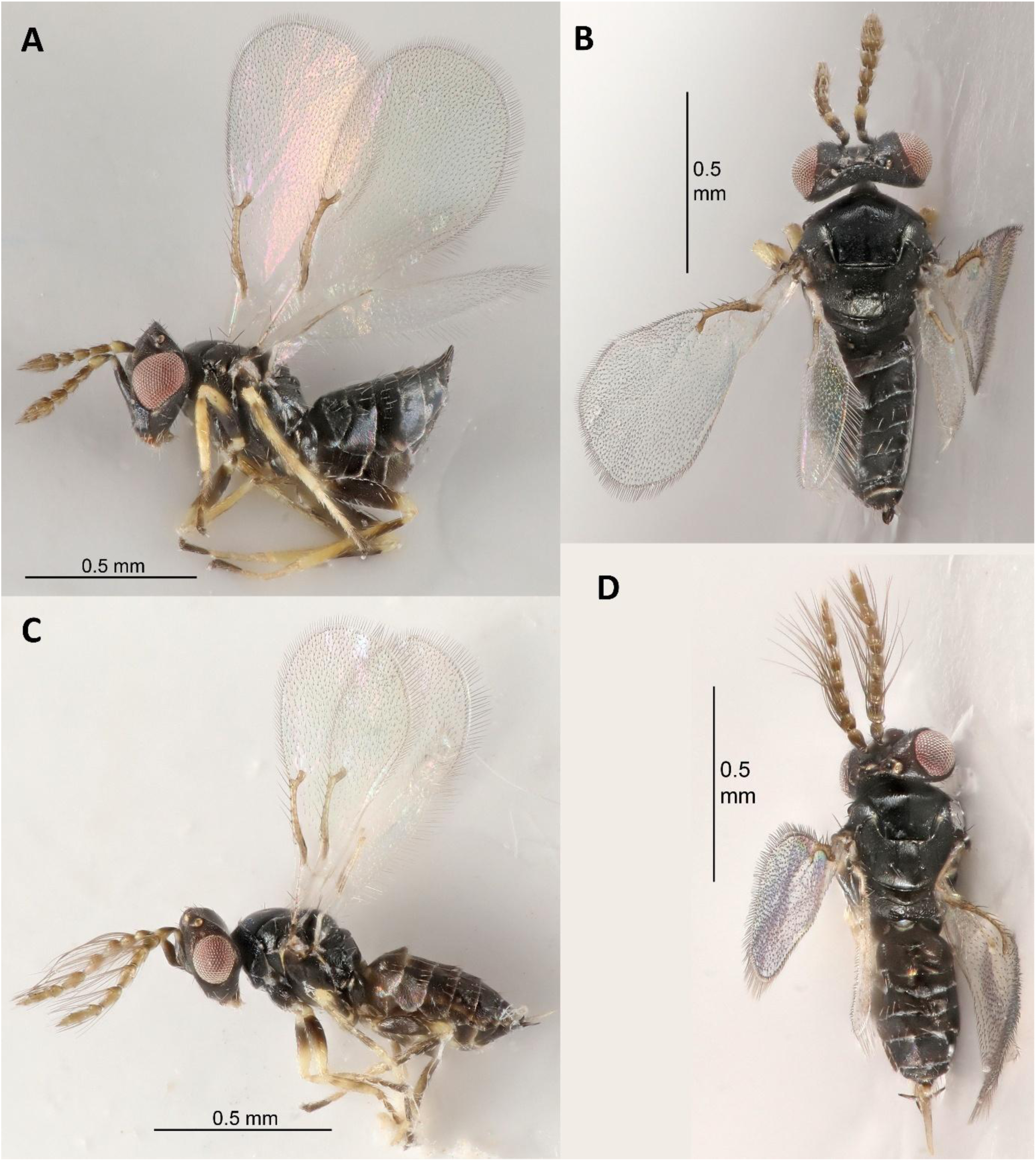
*Tamarixia citricola* Hansson and Guerrieri sp. nov. adult. (A) Female, lateral view, (B) Female, dorsal view, (C) Male, lateral view and (D) Male, dorsal view.

**ZooBank registration**: urn:lsid:zoobank.org:pub:26374181-2414-4C54-893B-0565F64F5A56

***Diagnosis*.** Fore wing (Figure 5A) with marginal vein short and thick, 5.4× as long as wide and only 0.16× as long as length of fore wing and 3.5× as long as stigmal vein; speculum relatively large, extending below marginal vein. Head, meso- and metasoma black, femora black with apex yellowish-white. Female antenna (Figure 5B) with F1 1.8× as long as wide and 0.8× as long as pedicel, F2 1.2× and F3 0.9× as long as wide; male antenna (Figure 5C) with plaque situated in basal ⅓; both sexes with scape black.

**Figure 5.**
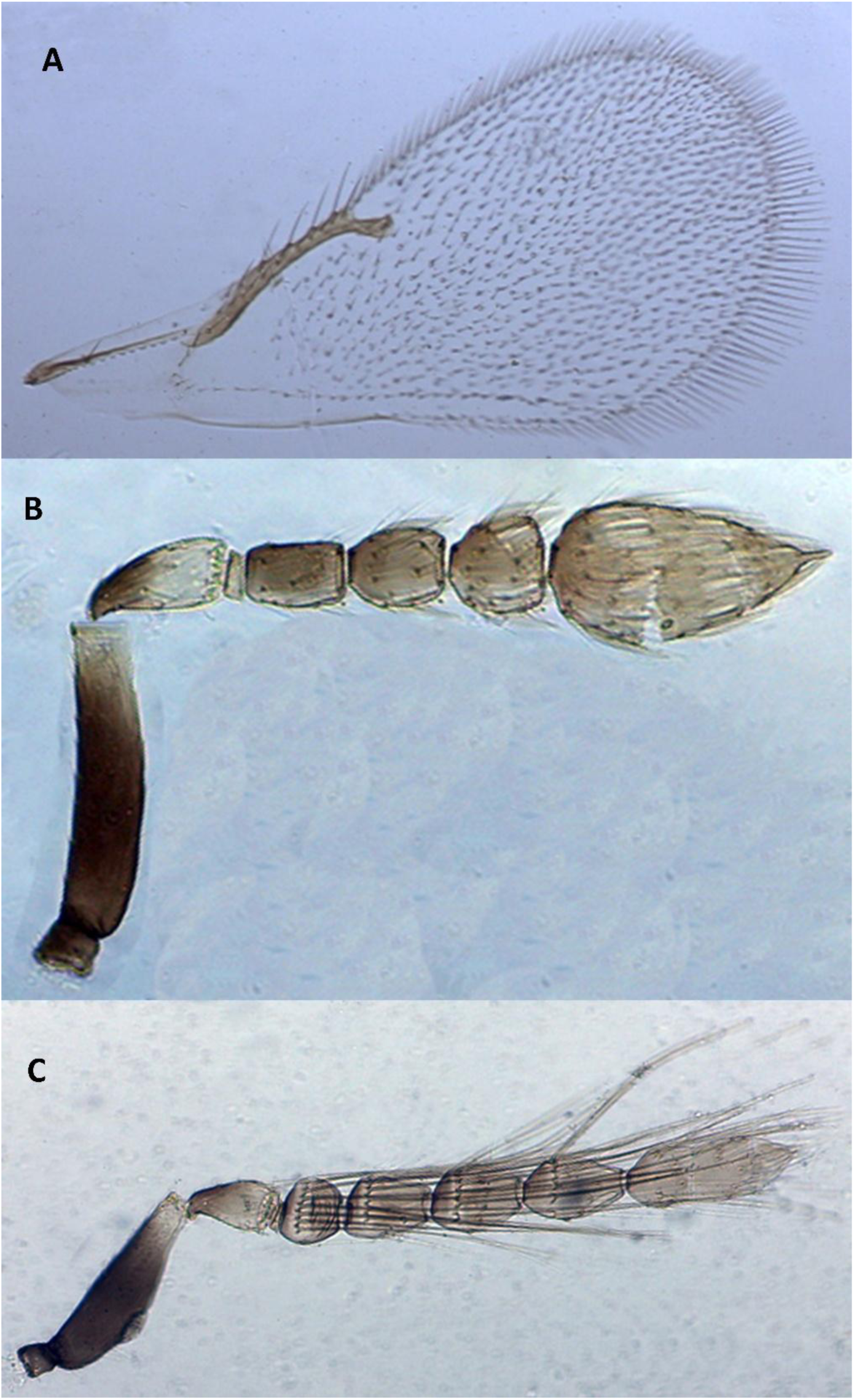
Diagnostic morphological traits of *Tamarixia citricola* Hansson and Guerrieri sp. nov. Female (A) Fore wing; (B) Antenna; Male (C) antenna.

***Female holotype***: length of body 1.0 mm.

Scape black with apico-ventral part white; pedicel with basal ½ black, apical ½ yellowish-white; flagellum dark brown. Head, meso- and metasoma black. Legs with coxae black; fore trochanter white with base black, mid and hind trochanters black; femora black with apex yellowish-white; fore and mid tibiae pale testaceous, hind tibia pale testaceous in basal ½, dark brown in apical ½; fore tarsus with T1–3 testaceous, T4 dark brown, mid and hind tarsi with T1–3 yellowish-white, T4 black. Wings hyaline, veins testaceous.

Midlobe of mesoscutum with weak reticulation, meshes elongate, with a complete but weak median groove. Mesoscutellum with distinctly weaker reticulation than mesoscutum and with smaller meshes; anterior setae as long as distance between submedian grooves and attached in the middle of mesoscutellum. Dorsellum with stronger reticulation than mesoscutum. Propodeum with weak reticulation, with a complete median carina.

Petiole very short and wide, a narrow stripe. Gaster with very weak reticulation throughout.

*Relative measurements*: head length, dorsal view 13; head length, frontal view 25; distance toruli to anterior ocellus 14; distance toruli to mouth margin 6; POL 9.5; OOL 3.5; lateral ocellus diameter 2; head width 29; mouth width 9; malar space 9; eye length 13.5; scape length 12.5; scape width 3; pedicel+flagellum length 27; pedicel length 5.5; pedicel width, dorsal view 2.5; F1 length 4.5; F1 width

2.5; F2 length 3.5; F2 width 3; F3 length 3.5; F3 width 4; clava length 10; clava width 5; C3 length 2.5; spicule length 1; mesosoma length 32; mesosoma width 29; midlobe of mesoscutum length 15; mesoscutellum length 12.5; mesoscutellum width 16; median part of mesoscutellum width (measured medially) 6; median part of mesoscutellum, width in anterior part 6; median part of mesoscutellum, width in posterior part 6; lateral part of mesoscutellum, width (measured medially) 4.5; dorsellum length 2.5; propodeum length 4.5; hind femur length 24; hind femur width 6.5; fore wing length 85; fore wing width 37; costal cell length 22; costal cell width (measured at widest part) 2; marginal vein length 14; stigmal vein length 4; gaster length 40; gaster width 23; Gt_7_ length (measured medially) 2; Gt_7_ width (measured at base) 2.5; longest cercal seta length 6.5; shortest cercal seta length 4.5; hypopygium length 23.

*Variation*. Length of body 0.9–1.1 mm. No additional variation in material examined.

***Male***. Length of body of body 0.8–1.0 mm.

Colour similar to female except with antennal flagellum pale testaceous to testaceous.

Antenna with scape widest in basal ⅓; plaque situated in lower ⅓; antennae with F1–F4 and C1 with a row of subbasal long setae. Otherwise as in female.

*Relative measurements*: head length, dorsal view 11; head length, frontal view 21; distance toruli to anterior ocellus 9.5; distance toruli to mouth margin 6; head width 26.5; mouth width 9.5; malar space 7.5; eye length 11; scape length 11; scape width 5; plaque length 2; distance apical part of plaque to apical part of scape 6; distance basal part of plaque to basal part of scape 3.5; pedicel length 4.5; pedicel+flagellum length 36; F1 length 3.5; F1 width 3; F2 length 5; F2 width 3.5; F3 length 5.5; F3 width 3; F4 length 5.5; F4 width 3; clava length 11; clava width 3; longest subbasal seta on F1 length 17; mesosoma length 29; mesosoma width 24; gaster length 34; gaster width 22.

**Hosts**. Parasitoid of Asian citrus psyllid (*Diaphorina citri* Kuwayama) (Hemiptera: Psyllidae).

**Distribution.** Cyprus.

**Material examined:** Holotype ♀ CYPRUS: Limassol, Eptagonia, 34°51’06.2”N 33°08’46.3”E, collected 30.x.2024 from the field, ex *Diaphorina citri* on *Citrus* sp. (emerged in cage 4.xi.2024) (deposited in Natural History Museum, London UK - NHMUK). Paratypes: 5♀ 5♂ CYPRUS: Nicosia, Peristona, 35°08’04.3”N 33°04’52.3”E, collected 31.x.2024 from the field, ex *Diaphorina citri* on *Citrus* sp. (emerged in cage 8.xi.2024) (deposited in NHMUK & Biological Museum (Entomology), Lund University, Lund, Sweden). Types 1♀ 3♂ *Tamarixia tremblayi* Domenichini, no collecting data, other than species name and « TYPE » on labels. (Domenichini Collection at Università Cattolica del Sacro Cuore, Piacenza, Italy).

## 4. Discussion

This study has identified an undescribed parasitoid species, *T. citricola* sp. nov., associated with nymphs of Asian citrus psyllid, *D. citri*, in Cyprus. This finding represents the first report of *T. citricola* parasitizing *D. citri* and the first record of this species in Europe. The area of origin for *T. citricola*, is at this time, assumed to be Cyprus.

Comparative analyses revealed clear genetic and morphological differences between *T. citricola* and the introduced species, *T. radiata*. DNA barcode sequences (COI) showed only 91–92% similarity between them, placing the field collected specimens from Cyprus in a separate clade in the phylogenetic analyses. This level of divergence is significant and consistent with interspecific differences within the genus (Om et al., 2017). Furthermore, distinct unambiguous diagnostic morphological characters differentiate *T. citricola* from *T. radiata*: in *T. citricola* gaster is completely black in both sexes, in *T. radiata* it is extensively yellow, more so in the female, in *T. citricola* femora and scape are predominantly black in both sexes, but whitish in *T. radiata*. In the key to European *Tamarixia* species by Graham (1991) both female and male *T. citricola* key out as *T. tremblayi* (Domenichini) (Domenichini, 1965), to which it is very similar. However, male *T. citricola* differ from that of *T. tremblayi* by the position of the sensorial plaque on the scape, situated in basal ⅓ of scape (in apical ⅓ in *tremblayi*, see fig. 327 in Graham [1991]). Females of *T. citricola* have black femora with a yellowish-white apex, gaster black, dorsellum 0.2× as long as mesoscutellum, and marginal vein 0.16× as long as length of fore wing. Conversely, in the *T*. *tremblayi* female, fore and mid femora are yellowish-white with basal ¼–⅓ dark brown, hind femur with basal ½ dark brown and apical ½ yellowish-white, gaster largelly yellow, dorsellum 0.3× as long as mesoscutellum, and marginal vein 0.13× as long as length of fore wing.

*Tamarixia radiata* is a solitary ectoparasitoid of *D. citri* nymphs, initially described from specimens reared from psyllids collected from lemons in Lyallpur, Punjab Pakistan in 1921 (Waterston, 1922) and widely used in classical biological control programs targeting *D. citri* (Étienne et al., 2001; Qureshi and Stansly, 2020; Hoddle and Pandey 2014). In contrast, the origin of *T. citricola* remains unclear. Its detection in orchards with no history of releases, suggests it may be a previously unrecognized native species, or possibly, accidentally introduced from some other unknown area. It is important to highlight that the parasitoid was not found during previous field observations carried out in the same orchards in 2023. To distinguish between these two competing possibilities, it will be crucial to identify psyllid host species (psyllid species native to Cyprus?) for *T. citricola* and to conduct further surveys in Cyprus and neighboring mainland countries for *T. citricola*. This work can be complemented with genetic studies to aid understanding of the area of origin, distribution patterns, and genetic population structures of *T. citricola* attacking *D. citri*. Regardless of the area of origin for *T. citricola*, currently available evidence confirms the coexistence in Cyprus of two eulophid species associated with *D. citri*: *T. radiata* (deliberately introduced) and *T. citricola* (unexpectedly detected, likely autochthonous, and a species new to science).

The simultaneous presence of *T. citricola* and *T. radiata* in the same agroecosystem raises important and interesting questions about their ecological interactions and implications for the biological control of *D. citri*. Although both are idiobiont ectoparasitoids that share the same host, they may differ in key ecological traits that facilitate their coexistence. Differences in thermal tolerance or seasonal activity may also exist. For example, one species may be better adapted to colder or warmer periods. Such temporal segregation could reduce direct competition. Additionally, assuming that *T. citricola* is native to Cyprus, this parasitoid may be opportunistically exploiting *D. citri*, a relatively new resource. As *T. radiata* populations increase in density and spread, interspecific competition for *D. citri* may result in the displacement of *T. citricola* and the host range of this parasitoid could regress to what it was originally before the invasion of *D. citri*.

In other systems where *T. radiata* competed with other parasitoid species of *D. citri* provide useful parallels for consideration. In Réunion, both *T. radiata* and the endoparasitoid *Diaphorencyrtus aligarhensis* (Shafee, Alam and Agarwal) (Hymenoptera: Encyrtidae) were introduced. *Tamarixia radiata* dominated, parasitizing ∼70% of nymphs compared to <20% by *D. aligarhensis* (Aubert, 1990, 1987). Similarly, in Florida, *T. radiata* established successfully following releases between 1999 and 2001, whereas *D. aligarhensis* failed to persist, likely due to interspecific competition and unfavorable environmental conditions (Rohrig et al., 2012). A similar outcome was observed in California, where *D. aligarhensis* failed to establish whilst *T. radiata* established readily and rapidly spread (Milosavljević et al., 2022).Laboratory studies suggested that *T. radiata* is more effective than *D. aligarhensis* at procuring and exploiting *D. citri* nymphs (Vankosky and Hoddle, 2019, 2017).

To understand the impact of *T. citricola* and *T. radiata* on *D. citri*, as well as their long-term population dynamics in Cyprus, it is essential to determine parasitism rates, host stage preferences, and whether these two parasitoid species exhibit behavioral discrimination against previously parasitized hosts. These parameters will help clarify whether the two parasitoid species are likely to complement each other or compete. Consistent differences could be expected in the interaction between these two congeneric species in respect to what has been so far reported for competing parasitoids of *D. citri*. If *T. citricola* provides functional complementarity, for example, being active when *T. radiata* is less effective for example, it may enhance biological control. Conversely, asymmetric competition could result in one species dominating and providing effective control. These possibilities warrant laboratory and field research.

From an integrated pest management (IPM) perspective, *T. citricola* holds considerable promise as a biological control agent targeting *D. citri* in the Mediterranean basin. The presence of *T. citricola* in Cypriot citrus orchards suggests good adaptation to local conditions, which could allow it to suppress *D. citri* populations throughout the year. Unlike exotic agents such as *T. radiata*, *T. citricola* is probably native and may already be present in other parts of the Mediterranean. If this possibility is confirmed, *T. citricola* could be promoted or redistributed for biological control of *D. citri* without facing regulatory constraints, a clear advantage when compared to importing and releasing non-native natural enemies, like *T. radiata*. However, to fully understand the potential of *T. citricola* for biological control of *D. citri*, the ecological role of this parasitoid needs to be better understood. In particular, the non-*D. citri* psyllids hosts (native psyllid species?) of *T. citricola* need to be identified. Host range and host specificity studies on *T. citricola* will be necessary too (see Hoddle and Pandey 2014; Bistline-East et al. 2015; Urbaneja-Bernat et al., 2019). These insights will guide how best to support the use of *T. citricola* against *D. citri* in the Mediterranean basin whether by conserving and enhancing natural populations or, where *D. citricola* is absent, by introducing this species into areas newly invaded by *D. citri*.

This study also highlights the critical role of integrative taxonomy (i.e., morphological and molecular analyses) in classical biological control of invasive pests. When combined with field surveys, molecular diagnostics and morphological analyses, enabled the detection of a second previously unknown parasitoid species where at the start of this classical biological control program, only one, *T. radiata*, was assumed to occur. Without this approach, *T. citricola* might have gone unnoticed or been misidentified as *T. radiata*, potentially leading to misinterpretations of field parasitism dynamics. In this case, an integrated multidisciplinary team approach has provided easy to use diagnostic tools for field entomologists and quarantine officers to distinguish *T. citricola* from *T. radiata*, an essential requirement for evaluating the identity, performance, distribution, and ecological roles of these biological control agents in Cyprus, and possibly other Mediterranean countries.

Finally, the presence of at least three *Tamarixia* species associated with quarantine-relevant psyllid species in Europe (*T. radiata* [*D. citri*], *T. dryi* [*T. erytreae*] and now *T. citricola* [*D. citri*]) illustrates the dynamic nature of biological control programs that are developed in response to invasive agricultural pests. Given the high invasion potential for *D. citri*, it is crucial to establish regional monitoring networks in the Eastern Mediterranean and the Middle East to detect early incursions of *D. citri* and to identify natural enemies associated with this pest if they exist. The invasion of *D. citri* into Cyprus illustrates both the challenges and opportunities emerging in the classical biological control of invasive pests. The discovery of a new parasitoid species during a classical biological control program highlights the importance of vigilance, integrative science, and international collaboration. As the Mediterranean citrus industry faces increasing HLB-related threats, identifying and integrating native and non-native natural enemy species, such as *T. citricola* and *T. radiata*, respectively, needs to be supported by sound pragmatic scientific principles. This approach will be key to building effective and sustainable control strategies for *D. citri* and the *C*. Liberibacter spp. this citrus pest vectors.

## Acknowledgments

The authors thank Prof. Emanuele Mazzoni, Università Cattolica del Sacro Cuore, Piacenza, Italy, for his help in locating and permission to loan the type of *Tamarixia tremblayi* (Domenichini). We thank María Antonia Gómez (IVIA) for her preliminary contribution to the morphological characterization.

## Statements & Declarations

## Author contributions

AU, AT, MPH, NS, and AMP conceived the study and initiated the first contacts with MSH and DJWM to launch the classical biological control (CBC) program in Cyprus. DJWM coordinated the shipment of *Tamarixia radiata* from California, and both DJWM and MSH provided background information essential for establishing the program. NS established the rearing of *Tamarixia radiata* in Cyprus and conducted its release. AU, AT, MPH, AMP, MS, DJWM, MSH, and NS participated in the design of parts of the study. AT, AMP, SG, MS, NS, CK, and LM contributed to field surveys, sampling, and data collection. MPH designed and ORR carried out the molecular taxonomy analyses. EG and CH performed the morphological characterization of parasitoid specimens, species identification, and species description. AU and EG wrote the first draft of the manuscript, which was further developed and completed by MPH and MSH. All authors read, reviewed, and approved the final version of the manuscript.

