## Supplementary material for "*Tamarixia citricola* Hansson and Guerrieri sp. nov. (Hymenoptera: Eulophidae): a new parasitoid of *Diaphorina citri* Kuwayamava (Hemiptera: Psyllidae) found during a classical biological control program in Cyprus": Supp Figure 1

**Supplementary Figure 1.** Comparison of the consensus COI gene fragment from a *Tamarixia radiata* laboratory-maintained colony in Cyprus, with publicly available COI sequences of *T. radiata* retrieved from GenBank.

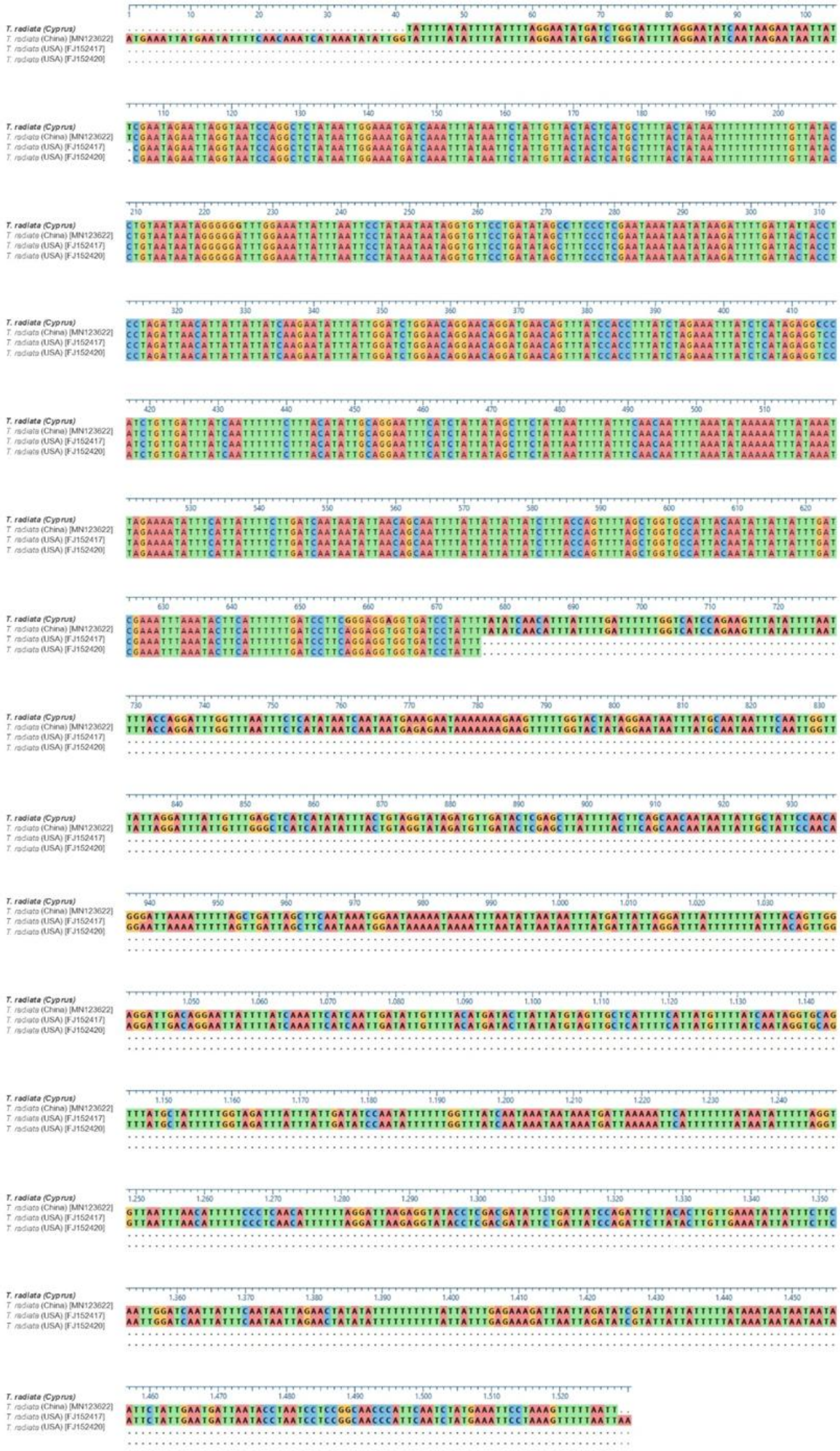
T. radiata (Cyprus)
T. radiata (China) [MN123622]
T. radiata (USA) [FJ152417]
T. radiata (USA) [FJ152420]
